# Metabolic Reprogramming and Therapeutic Vulnerabilities in the Tumor Microenvironment Revealed by Multi-scale Network Geometry

**DOI:** 10.1101/2025.11.16.688737

**Authors:** Handan Cetin, Paul Bogdan, Shannon M Mumenthaler, Stacey D Finley

## Abstract

The tumor microenvironment comprises diverse cell populations that coordinate metabolic activities to sustain malignant growth, yet the systems-level organization of these interactions remains poorly understood. Here, we present an integrated framework combining single-cell transcriptomics, genome-scale metabolic modeling, and multi-scale network geometry to decode metabolic coordination in colorectal cancer. We demonstrate that FAP**^+^** cancer-associated fibroblasts and MARCO**^+^** tumor-associated macrophages undergo extensive reprogramming, establishing metabolic division of labor: fibroblasts specialize in amino acid and fatty acid metabolism while macrophages adopt cancer-like nucleotide biosynthesis programs. Systematic knockout analysis identified 19 tumor-selective vulnerabilities in branched-chain amino acid catabolism, with MAOB validated as a prognostic marker through patient survival analysis. To reveal architectural organization, we applied multifractal geometric characterization and Ollivier-Ricci curvature analysis for the first time to flux-weighted metabolic networks derived from context-specific genome-scale models. While conventional network metrics failed to distinguish tumor from normal phenotypes, multifractal analysis successfully separated tissue states through coordinated architectural changes across hierarchical scales. Role transition analysis revealed that 20–25% of metabolites undergo functional reorganization, with prostaglandin and bile acid derivatives emerging as critical communication hubs between stromal populations. Curvature analysis identified pathway-specific geometric remodeling in fatty acid metabolism (fibroblasts) and leukotriene metabolism (macrophages). Our findings establish that metabolic adaptation represents ecosystem-level network reorganization rather than isolated pathway changes, providing a generalizable framework for identifying therapeutic strategies targeting cooperative metabolic networks.

## 1 Introduction

Colorectal cancer (CRC) remains one of the most prevalent and lethal malignancies worldwide, ranking third in cancer incidence and second in mortality [1, 2]. Despite advances in surgical techniques and targeted therapies, the five-year survival rate for metastatic disease remains below 15% [3]. This therapeutic challenge stems partly from our incomplete understanding of how diverse cell populations within the tumor microenvironment (TME) coordinate their metabolic activities to sustain tumor growth and evade therapeutic interventions [4]. Unfortunately, current cancer treatments often fail because they target cancer cells in isolation, with little understanding of the impact on the complex ecosystem of supporting cells that enable tumor growth.

The TME comprises a complex ecosystem where cancer cells coexist with various stromal populations including fibroblasts, immune cells, endothelial cells, and other supporting cell types [5, 6]. Rather than serving as passive bystanders, these stromal cells actively participate in tumor progression through metabolic crosstalk, creating what has been termed a “metabolic ecosystem” [7, 8]. Cancer-associated fibroblasts (CAFs) represent the most abundant stromal component in many solid tumors [9–11], while tumor-associated macrophages (TAMs) comprise up to 50% of the tumor mass in some cancers [12, 13].

Recent single-cell RNA sequencing studies have revealed unprecedented cellular heterogeneity within colorectal tumors [14–16], including the presence of distinct subsets of CAFs and TAMs. Of note, Qi et al. identified tumor-specific FAP^+^ fibroblasts and MARCO^+^ (SPP1^+^) macrophages that were highly correlated across multiple patient cohorts and associated with poor prognosis [17]. However, these studies primarily focused on transcriptional profiles without systematically investigating the metabolic dimensions of stromal heterogeneity.

Metabolism represents a fundamental cellular process that both responds to and shapes the TME [18]. Emerging evidence suggests that metabolic reprogramming extends beyond cancer cells to encompass the entire tumor ecosystem [19]. CAFs have been shown to undergo reverse Warburg metabolism, producing lactate and other metabolites that fuel cancer cell growth [20]. TAMs adopt distinct metabolic programs depending on their polarization state, with M1-like macrophages relying on glycolysis while M2-like macrophages utilize oxidative phosphorylation [21].

The metabolic interplay between different cell types creates emergent properties that cannot be understood by studying individual populations in isolation. Examples include CAFs providing alanine to cancer cells in pancreatic cancer [22], glutamine-lactate metabolic cycling in ovarian cancer [23], and fatty acid transfer by adipocytes in breast cancer [24]. These examples highlight the importance of understanding metabolism at the ecosystem level.

Several computational approaches exist to investigate the metabolic features of cells and how those features influence cells in the TME at the systems-level, including genome-scale metabolic models and network analysis. Such tools can be applied to provide quantitative and mechanistic insights into the metabolic ecosystems of CRC. Genome-scale metabolic models (GEMs) integrate multi-omics data to predict metabolic fluxes and identify vulnerabilities [25, 26]. Context-specific model reconstruction algorithms enable generation of cell type-specific GEMs that capture metabolic diversity within tissues [27]. While traditionally analyzed using linear optimization for a single objective such as growth, recent advances enable exploration of alternative metabolic states. In particular, random flux sampling allows unbiased exploration of the solution space, revealing metabolic capabilities beyond growth optimization [28].

Biological network analysis provides powerful tools for understanding metabolic organization beyond individual pathways. Metabolic networks exhibit scale-free topology, where a few highly connected metabolites (hubs) dominate network structure [29]. This architecture creates both robustness and vulnerability: deletion of most nodes has minimal impact, but targeting hubs can collapse network function [30]. Network medicine has successfully identified drug targets by analyzing topology [31], but most analyses treat networks as static entities, missing how architecture adapts to different conditions.

Geometric analysis of networks reveals organizational principles invisible to traditional topological metrics [32]. Multifractal analysis quantifies complexity across scales [33], while Ollivier-Ricci curvature distinguishes tightly integrated modules from sparse regions [34]. These geometric properties have been linked to network function in protein interaction networks [35] and brain networks [36], but remain unexplored in metabolic networks.

Here, we present an integrated systems biology approach to decode metabolic coordination in the CRC microenvironment. By combining genome-scale metabolic modeling with advanced network topology and geometric analyses, we reveal how FAP^+^ fibroblasts and MARCO^+^ macrophages undergo coordinated metabolic reprogramming that extends beyond nutrient exchange to encompass fundamental reorganization of metabolic network architecture, compared to normal tissue. We demonstrate that these populations adopt complementary metabolic strategies creating a metabolic division of labor that sustains tumor growth. Through novel application of metabolite role transition analysis, multifractal characterization, and curvature-based geometric analysis, we identify emergent metabolic vulnerabilities that cannot be identified through traditional approaches such as flux balance analysis. Our findings reveal that metabolic adaptation in CRC involves coordinated architectural reorganization across multiple scales, providing a road map for developing and implementing therapeutic strategies targeting the metabolic ecosystem. Our results establish a quantitative framework for decoding ecosystem-level metabolic organization in the TME and is applicable across cancers. This work enables identification of therapeutic strategies that intervene in cooperative metabolic networks instead of single molecular pathways.

## 2 Results

### 2.1 Context-specific metabolic models reveal preserved cell identity and extensive stromal reprogramming

Tumor ecosystems coordinate metabolic reprogramming across diverse cell populations, yet the extent of this coordination in tumors remains poorly understood. To systematically map metabolic relationships in the CRC ecosystem, we re-analyzed single-cell RNA sequencing data comprising 54,103 cells from CRC patients obtained from Qi et al. (2022). This dataset included cells from tumor tissue and tumor-adjacent normal tissue, allowing direct comparison of metabolic states in matched conditions. Building on their cell type annotations, we generated context-specific genome-scale metabolic models (GEMs) — computational frameworks that integrate gene expression data with biochemical knowledge to reconstruct cell type-specific metabolic networks. We used established algorithms to investigate metabolic cooperation between tumor-enriched populations. In addition to representing the full network of a cell’s metabolism, the GEM also includes a biomass reaction that represents cell growth.

While we present results for all fibroblast and macrophage subtypes identified by Qi et al. to be complete, we particularly focused our analysis on two stromal populations that Qi et al. identified as strongly correlated across patient cohorts and associated with poor prognosis: FAP^+^ fibroblasts and MARCO^+^ (SPP1^+^) macrophages. These populations represent key tumor-associated stromal cells that are highly enriched in tumor tissue compared to adjacent normal tissue (Figure 1A), making them ideal candidates for understanding how stromal cells adapt metabolically to support tumor growth. We note that malignant epithelial cells were detected in both tumor and normal tissue contexts, reflecting the field cancerization phenomenon [37] where tumor-adjacent tissue harbors pre-malignant or early malignant cells. The presence of malignant epithelial cells in both conditions enabled us to use these cells as a reference point for cancer-associated metabolic programs when comparing stromal cell adaptations.

**Fig. 1:**
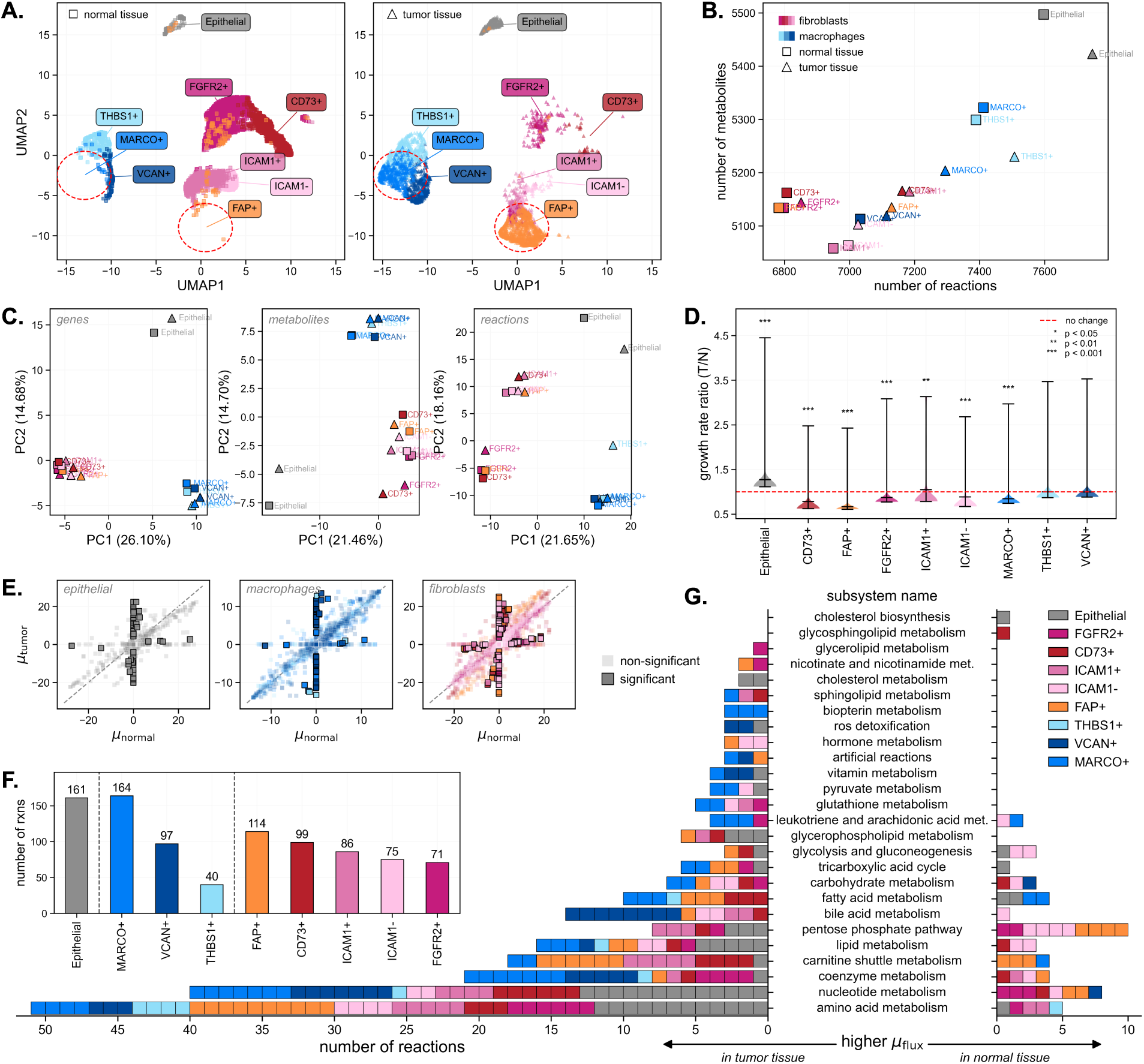
Context-specific metabolic models reveal metabolic reprogramming in fibroblasts and macrophages in the tumor microenvironment. **A.** UMAP visualization showing cellular abundance in tumor-adjacent normal and tumor tissue, demonstrating enrichment of FAP^+^ fibroblasts and MARCO^+^ macrophages in tumor conditions. **B.** Metabolic model complexity showing relationship between reactions and metabolites for each context-specific model. **C.** PCA of binary gene, metabolite, and reaction vectors showing cell type clustering across all populations, fibroblast subtypes, and macrophage subtypes, confirming metabolic identity preservation regardless of tissue origin. **D.** Growth rate fold changes showing decreased biomass production in tumor-associated cells compared to normal counterparts. Dashed red line shows a ratio of 1, indicating the same growth rate for tumor and normal conditions. **E.** Scatter plots of mean reaction fluxes comparing tumor versus normal conditions for fibroblasts and macrophages. **F.** Quantification showing number of significantly differentially active reactions per cell type. **G.** Pathway-level distribution of differentially active reactions across metabolic subsystems for each cell type.

Context-specific GEMs revealed architectural diversity in metabolic networks (Figure 1B). Epithelial malignant cells maintained the largest networks with over 5,400 metabolites and 7,500 reactions, while fibroblasts showed the smallest networks, and macrophages exhibited intermediate network size. This architectural diversity raised a fundamental question: do tumor conditions rewire the core metabolic identity of cells? Hierarchical clustering of reaction presence revealed that metabolic network architecture serves as a robust cellular fingerprint, preserving lineage identity even under tumor stress. Cells clustered strictly by lineage rather than by tumor versus normal origin (Supplementary Figure S1). Principal component analysis of binary gene, metabolite, and reaction vectors confirmed clear separation between cell types regardless of tissue origin (Figure 1C).

To quantify functional metabolic differences, we performed differential flux analysis using random sampling of context-specific GEMs. All cell types except malignant epithelial cells showed decreased growth rates in their tumor counterparts, compared to normal (Figure 1D), suggesting stromal cells adopt metabolic strategies that prioritize tumor-supporting functions over self-proliferation.

Across all cell subtypes, nearly all of the reactions with significantly different fluxes showed higher activity in tumor conditions (Figure 1E). MARCO^+^ macrophages exhibited the most extensive metabolic reprogramming with 164 significantly differentially active reactions, followed by malignant epithelial cells (161 reactions) and FAP^+^ fibroblasts (114 reactions) (Figure 1F).

### 2.2 Metabolic division of labor: FAP^+^ fibroblasts specialize in metabolic supply while MARCO^+^ macrophages adopt cancer-like biosynthetic programs

While the quantitative analysis demonstrated extensive metabolic reprogramming in FAP^+^ fibroblasts and MARCO^+^ macrophages, the functional consequences of these changes remained unclear. To understand how metabolic rewiring translates into tumor-supporting capabilities, we performed pathway-level analysis using metabolic subsystem annotations. Detailed reaction-level changes for all metabolic subsystems are shown in Supplementary Figure S2.

Pathway-level analysis revealed the functional significance of metabolic changes across different subsystems (Figure 1G). FAP^+^ fibroblasts showed extensive reprogramming in amino acid metabolism (10 upregulated reactions) and carnitine shuttle metabolism (9 reactions, 6 upregulated), suggesting enhanced capacity for protein synthesis and fatty acid oxidation. Notably, compared to normal conditions, tumor FAP^+^ fibroblasts showed downregulation in the pentose phosphate pathway and nucleotide metabolism, contrasting with upregulation of these pathways in malignant epithelial cells.

MARCO^+^ macrophages demonstrated distinct metabolic profiles, with the most changes in coenzyme and nucleotide metabolism (7 reactions each, all upregulated). This reflects high metabolic demands for immune effector functions and potential nucleotide provision to rapidly proliferating cancer cells. The changes in paralleled patterns observed in malignant epithelial cells (13 upregulated nucleotide reactions). MARCO^+^ macrophages also showed significant alterations in leukotriene and arachidonic acid metabolism, indicating reprogramming of inflammatory lipid mediator pathways.

Both FAP^+^ fibroblasts and MARCO^+^ macrophages showed substantial upregulation of amino acid metabolism in tumor conditions, paralleling malignant epithelial cells and indicating the central importance of amino acid availability for tumor ecosystem function. However, the populations diverged in their approach to biosynthetic metabolism. Malignant epithelial cells and MARCO^+^ macrophages both exhibited cancer-like metabolic reprogramming, with increased nucleotide metabolism and pentose phosphate pathway activity. In contrast, FAP^+^ fibroblasts displayed a complementary phenotype, showing decreased nucleotide metabolism while upregulating fatty acid oxidation and amino acid production pathways. Altogether, the changes reveal a metabolic division of labor between FAP^+^ fibroblasts and MARCO^+^ macrophages.

### 2.3 Computational gene knockout analysis identifies selective vulnerabilities in tumor-associated cell populations with clinical prognostic value

Building on our context-specific metabolic models, we performed systematic knockout analysis to identify and validate therapeutic targets within the metabolic network. Targets were filtered using dual criteria designed to maximize the therapeutic window: (1) mean knockout effect across tumor cell types → 0.4 (indicating substantial growth inhibition), and (2) mean knockout effect across normal cell types ↑ 0.6 (ensuring minimal toxicity to healthy tissue) (Figure 2A).

**Fig. 2:**
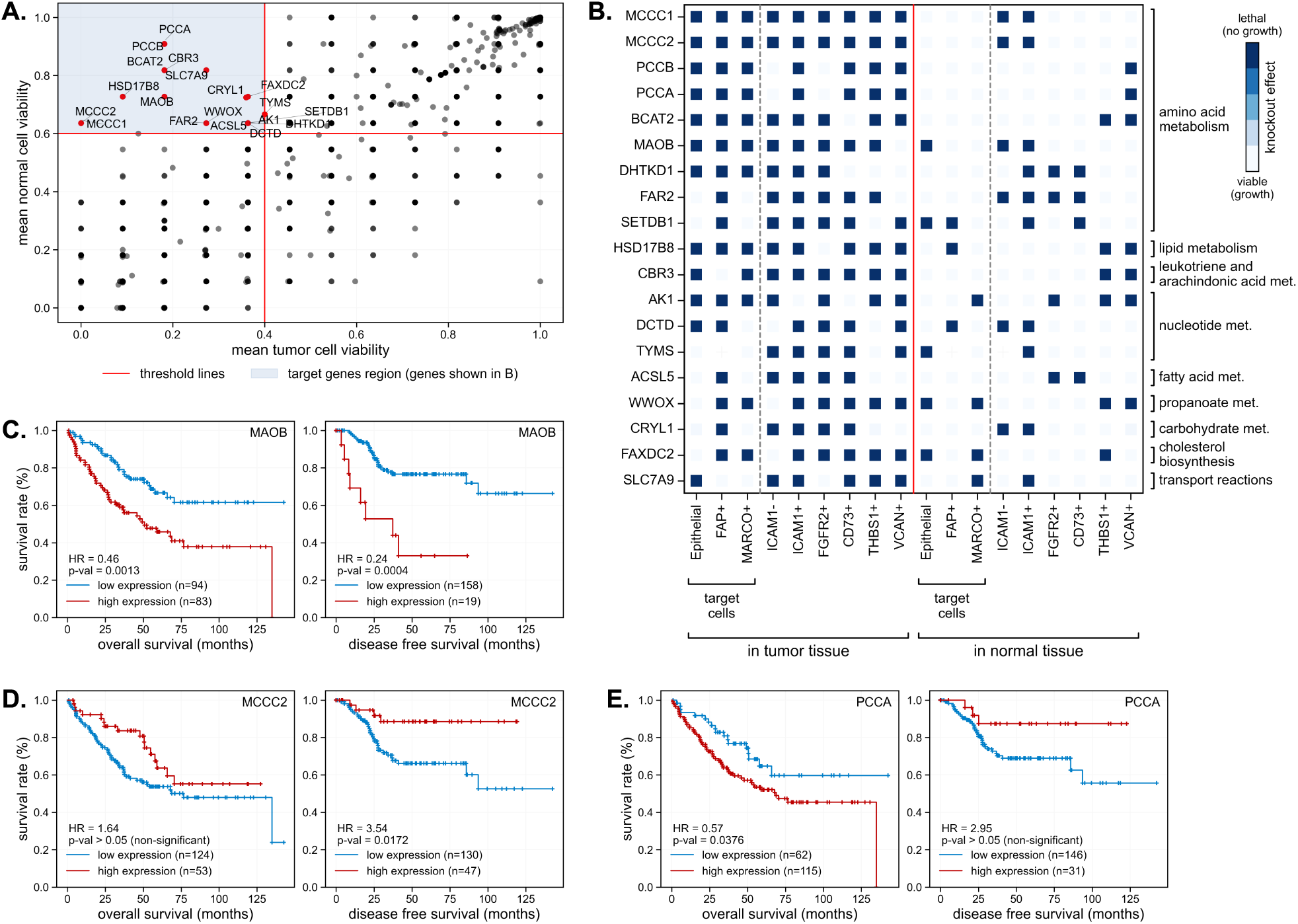
Computational gene knockout analysis identifies metabolic vulnerabilities with clinical prognostic value. **A.** Scatter plot showing the relationship between mean cell viability in tumor versus normal tissues for all knockout targets. Points in the target region (blue shaded area) represent genes with selective toxicity to tumor cells while sparing normal cells. **B.** Heatmap showing knockout ratios for candidate metabolic targets across tumor-associated cell populations (malignant epithelial cells, FAP^+^ fibroblasts, MARCO^+^ macrophages) and their normal tissue counterparts. Blue intensity indicates essentiality (knockout ratio = 1.0 = dark blue represents complete lethality). Genes are grouped by metabolic pathway annotations. **C.** Kaplan-Meier survival curves for three of top targets (MCCC2, PCCA, MAOB) showing both overall survival (left panels) and disease-free survival (right panels) in the GSE17536 colorectal cancer cohort.

This systematic approach applied across all metabolic genes, identified 19 candidate genes with optimal therapeutic windows (Figure 2B). MCCC2 and MCCC1, encoding 3-methylcrotonyl-CoA carboxylase subunits involved in leucine catabolism, emerged as the most selective candidates, demonstrating complete lethality across all tumor-associated populations while sparing normal counterparts. This exceptional selectivity validates our pathway analysis showing upregulated amino acid metabolism as a shared vulnerability across tumor populations. Following closely, BCAT2 (branched-chain amino acid transaminase 2) and PCCA and PCCB (propionyl-CoA carboxylase subunits). Additionally, MAOB (monoamine oxidase B) exhibited similarly robust tumor-selective lethality, further emphasizing the critical dependence on branched-chain amino acid catabolism in tumor-associated cells.

To validate clinical relevance of our computational predictions, we examined patient survival correlations using the GSE17536 dataset. Among the top candidates, we selected three genes showing differential prognostic associations (Figure 2C). Most promising, MAOB high expression consistently predicted poor outcomes in both overall survival and disease-free survival, establishing it as a robust biomarker of aggressive disease. MCCC2 demonstrated context-dependent survival correlations, with high expression predicting worse overall survival but improved disease-free survival. This suggests the gene has dual roles in treatment response versus long-term disease control. High expression of PCCA correlated with improved overall survival without affecting disease-free survival, indicating selective impact on metastatic progression.

These contrasting patterns demonstrate that metabolic dependencies manifest differently across disease stages, and that FBA-based knockout analysis, while valuable for initial screening, assumes biomass maximization that may not capture the complex selective pressures operating in clinical contexts due to the complex tumor ecosystem. The divergent survival associations indicate that effective therapeutic targeting requires understanding metabolism as an integrated network system rather than isolated gene perturbations.

### 2.4 Network architecture maintains cell identity while enabling tumor-specific adaptations

The metabolic division of labor and selective therapeutic vulnerabilities we identified through genomescale modeling arise due to complex metabolic networks whose organizational principles remain unclear. To understand how cell type-specific metabolic reprogramming translates into network-level organizational changes, we constructed directed metabolite-centric graphs from each cell type-specific GEM, where nodes represent metabolites and directed edges represent metabolic reactions connecting substrates to products with edge weights assigned as mean sampled flux values. We employed six established centrality measures that capture different aspects of metabolite importance (Figure 3A): degree centrality identifying highly connected hubs, betweenness centrality revealing critical bottlenecks between network modules, closeness centrality measuring network-wide accessibility, PageRank assessing recursive importance through connections to other important nodes, load centrality quantifying traffic burden, and eigenvector centrality indicating structural prominence. An example illustrating each centrality metric is also provided, where emphasis is placed on the biological example (colored node), rather than interpretation of the nodes to which it is connected. Together, these centrality measures enable comprehensive characterization of how individual metabolites function within the broader metabolic network and how their roles change between normal and tumor conditions (Figure 3B).

**Fig. 3:**
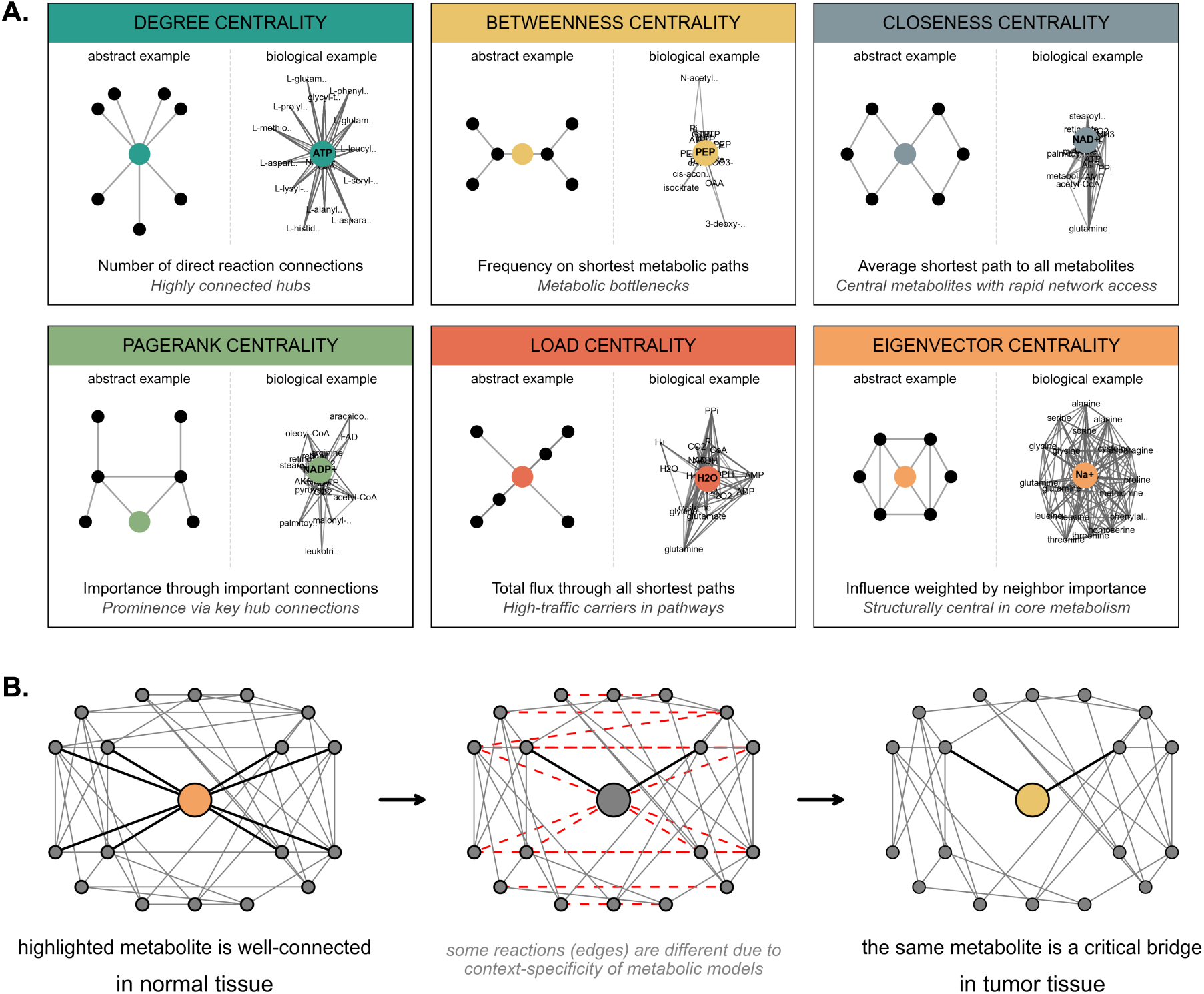
Network centrality measures and their biological interpretations in metabolic networks **A.** Six centrality metrics identify distinct functional roles: degree centrality (highly connected hubs), betweenness centrality (metabolic bottlenecks), closeness centrality (central metabolites with rapid network access), PageRank centrality (prominence via key hub connections), load centrality (high-traffic carriers in pathways), and eigenvector centrality (structurally central in core metabolism). Abstract network representations illustrate each measure alongside biological examples from metabolic networks. **B.** Context-specific metabolic network rewiring showing the same metabolite (highlighted node) exhibits distinct network topologies when analyzed in different biological contexts (in normal vs. tumor tissue).

Simply considering the sizes of the graphs for each cell type does not show differences between cell types (Figure 4A). However, analysis of fundamental network properties revealed distinct organizational adaptations reflecting cellular metabolic complexity and functional demands (Figure 4B). Network density, referring to the connectivity of the metabolic nodes, demonstrated variable patterns across cell types, with macrophage populations showing increased density under tumor conditions. Average edge weights, corresponding to mean reaction fluxes, revealed opposing trends between the two stromal populations: macrophages showed decreased flux magnitudes while fibroblasts demonstrated increased values. In contrast, degree heterogeneity (a measure of connectivity variance across metabolites), maximum PageRank (indicating the most influential metabolic hub based on recursive importance), and balanced nodes (representing metabolites with similar numbers of incoming and outgoing connections) remained relatively stable between normal and tumor conditions across cell types. Statistical analysis revealed that these apparent differences did not reach significance when comparing tumor versus normal conditions at single-scale network measures.

**Fig. 4:**
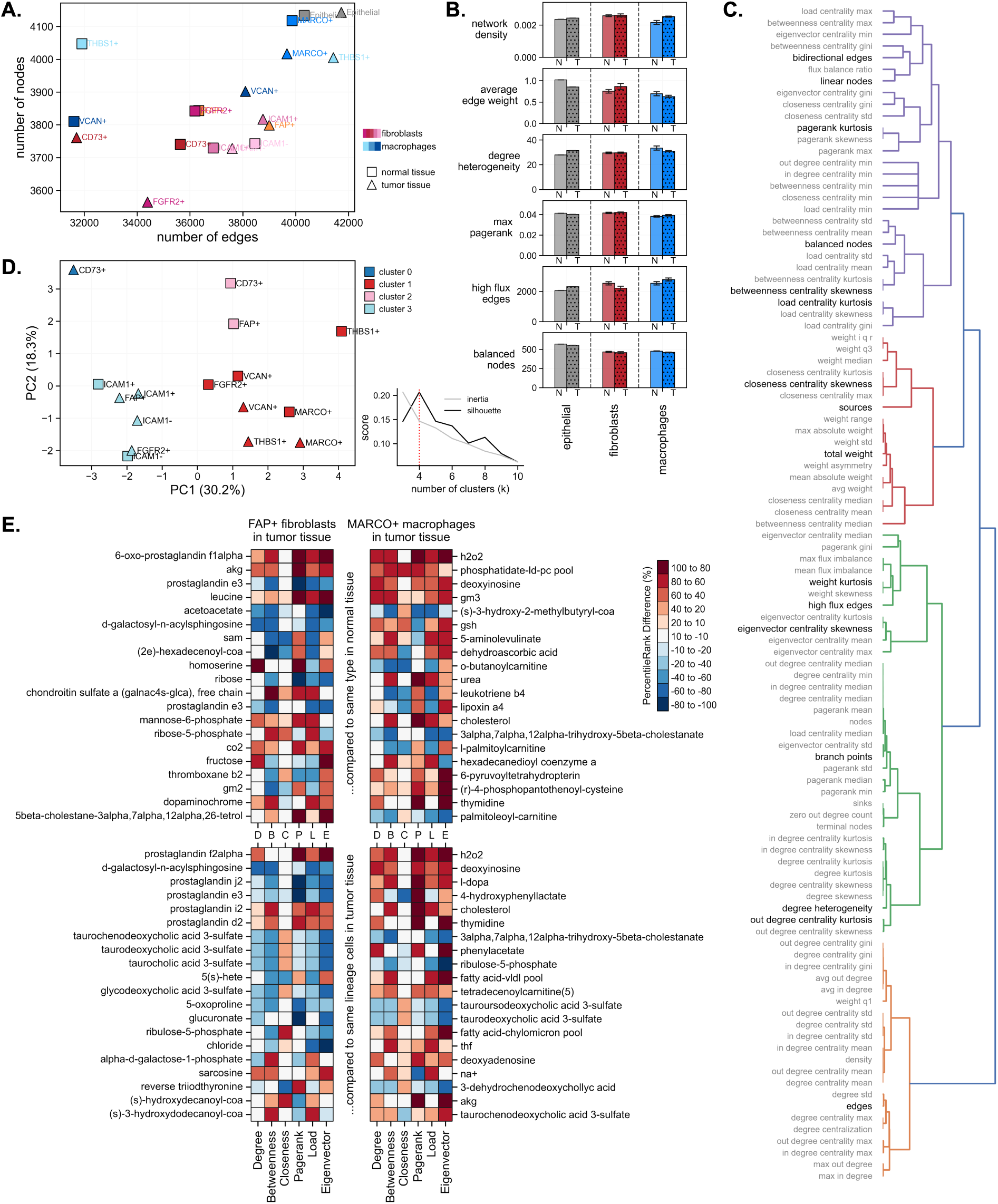
Network topology analysis reveals cell type-specific metabolic reorganization patterns and identifies key metabolites driving tumor-associated adaptations. **A.** Network size comparison showing number of nodes (metabolites) versus edges (reactions) for each cell type-specific model. **B.** Network properties showing density, average weight, degree heterogeneity, maximum PageRank, high-flux edges, and balanced nodes proportion. **C.** Dendrogram of 101 calculated topological metrics, highlighting 16 independent features (shown in black), with correlation matrix of final features confirming independence. **D.** K-means clustering of networks using independent topological features, with silhouette and inertia analysis shown confirming optimal cluster number of 4. **E.** Metabolite centrality rank difference heatmaps for FAP+ fibroblasts (left) and MARCO+ macrophages (right). Top panels show tumor versus normal tissue comparisons identifying tumor-specific metabolic adaptations. Bottom panels show comparisons against other cell types within the same lineage in tumor tissue. Positive values (red) indicate increased centrality in the target population, negative values (blue) indicate decreased centrality.

To systematically characterize network topology beyond these basic network features, we calculated 101 quantitative metrics encompassing structural properties, centrality measures, and connectivity patterns (Figure 4C). Feature selection through variance filtering reduced this to 45 informative metrics, followed by correlation filtering to identify 16 independent features that capture the essential topological differences between networks. Importantly, none of these topological features showed clear separation between tumor and normal conditions when analyzed individually (Supplementary Figure S3), indicating that metabolic network reorganization involves subtle, coordinated changes across multiple architectural dimensions rather than dramatic alterations in single network properties.

Unsupervised clustering analysis using these 16 key topological features revealed four distinct network organization patterns (Figure 4D), with optimal cluster number determined through silhouette analysis and inertia evaluation. MARCO^+^ macrophages clustered with other macrophage populations (cluster 1), demonstrating conservation of lineage-specific network organization principles. Most intriguingly, FAP^+^ tumor fibroblasts clustered with ICAM1 telocytes and FGFR2^+^ fibroblasts (cluster 3), providing compelling evidence that FAP^+^ fibroblasts represent activated states of these precursor populations rather than fundamentally distinct cell types. This clustering pattern supports the hypothesis proposed by Qi et al. that tumor-specific FAP^+^ fibroblasts likely originate from FGFR2^+^ fibroblasts or ICAM1^+^ telocytes through phenotypic transition.

### 2.5 Prostaglandin and bile acid metabolism emerge as central hubs in stromal metabolic networks

While the network-level topological analysis revealed subtle coordinated changes that did not achieve statistical significance at single scales, we next examined whether tumor-normal differences might be detectable at the individual metabolite level. To identify specific metabolites that drive the network reorganization patterns observed in FAP^+^ fibroblasts and MARCO^+^ macrophages, we focused on the centrality metrics introduced and defined in Figure 3 to calculate metabolite centrality rank differences (Figure 4E).

Analysis of FAP^+^ fibroblasts between tumor and normal conditions revealed dramatic reorganization of lipid mediator networks. The most prominently reorganized metabolites included prostaglandin derivatives (6-oxo-prostaglandin f1alpha and prostaglandin e3), showing consistently elevated centrality ranks across multiple measures in tumor conditions. Fatty acid metabolites and amino acids also demonstrated increased centrality, while sphingolipid intermediates and one-carbon metabolism metabolites showed decreased centrality in tumor conditions.

Comparison of FAP^+^ tumor fibroblasts against other tumor-associated fibroblast subtypes revealed their unique metabolic specialization within the stromal compartment. FAP^+^ fibroblasts showed distinctive elevation in prostaglandin metabolism (prostaglandin f2alpha, prostaglandin j2, prostaglandin i2, and prostaglandin d2) when compared to other fibroblast populations. Additionally, bile acid metabolites including taurochenodeoxycholic acid 3-sulfate, taurodeoxycholic acid 3-sulfate, and taurocholic acid 3-sulfate demonstrated decreased centrality specifically in FAP^+^ fibroblasts.

MARCO^+^ macrophage analysis between tumor and normal conditions demonstrated coordinated reorganization of nucleotide and lipid metabolism networks. Key metabolites showing elevated centrality in tumor conditions included nucleotide derivatives (h2o2, deoxyinosine, gm3), cholesterol-related compounds, and phospholipid metabolites. The prominence of deoxyinosine and related nucleotide salvage intermediates indicates that MARCO^+^ macrophages acquire enhanced capacity for nucleotide recycling in tumor conditions.

Comparison of MARCO^+^ tumor macrophages against other cell subtypes revealed their specialized metabolic niche within the immune compartment. MARCO^+^ macrophages demonstrated increased centrality in fatty acid metabolism intermediates and specific prostaglandin derivatives, compared to other tumor-associated macrophage subtypes. Notably, bile acid metabolites showed increased centrality in MARCO^+^ macrophages, paralleling the pattern observed in FAP^+^ fibroblasts and suggesting potential metabolic cooperation between these populations.

### 2.6 Metabolite role transitions reveal potential therapeutic bottlenecks and communication networks

We analyzed metabolite role transitions by classifying each metabolite according to its dominant centrality measure and tracking changes between normal and tumor conditions. Using the six centrality measures described above, we classified each metabolite into distinct functional roles based on their highest-ranking centrality: hubs (degree-dominant), bottlenecks (betweenness-dominant), connectors (closeness-dominant), influential nodes (PageRank-dominant), traffic carriers (load-dominant), and structural centers (eigenvector-dominant) (Figure 5A).

**Fig. 5:**
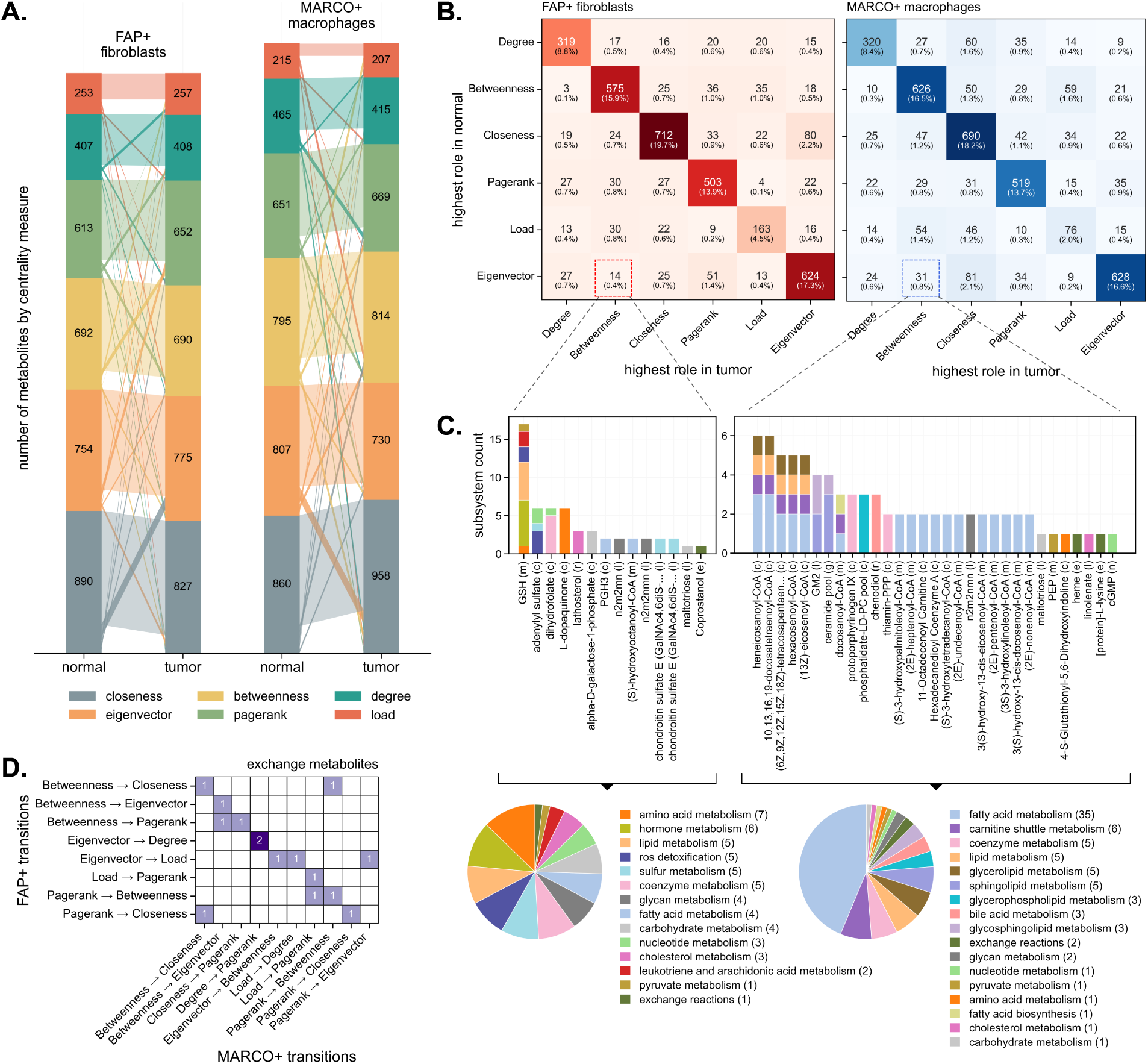
Metabolite role transitions reveal therapeutic targets and intercellular communication networks. **A.** Alluvial diagrams showing role transitions between normal and tumor conditions for FAP+ fibroblasts (left) and MARCO+ macrophages (right). Flow width represents the number of metabolites transitioning between roles, with colors indicating the six functional categories. **B.** Transition matrices quantifying role changes for FAP+ fibroblasts (left, 3,609 metabolites) and MARCO+ macrophages (right, 3,793 metabolites). **C.** Subsystem analysis of eigenvector-to-betweenness transition metabolites, showing metabolic pathways that create new bottleneck control points. Pie charts display the distribution of transition metabolites across metabolic subsystems for FAP+ fibroblasts (left) and MARCO+ macrophages (right). Bar plots show the number of metabolites in each subsystem category. **D.** Analysis of extracellular metabolite role transitions revealing intercellular communication networks.

Analysis revealed extensive metabolic reorganization in both FAP^+^ fibroblasts and MARCO^+^ macrophages (Figure 5B). In FAP^+^ fibroblasts, 3,609 metabolites were tracked, with 80.2% maintaining their original functional roles while 19.8% underwent role transitions (Figure 5C, left panel). Among the transitions, the most prominent change involved closeness-to-eigenvector shifts (80 metabolites, 2.2%), where metabolites providing efficient network-wide connectivity in normal tissue evolved to occupy structurally central positions in the TME. The second most significant transition was eigenvector-to-PageRank (51 metabolites, 1.4%), indicating that structurally central metabolites in normal tissue gained prominence through connections to other important nodes rather than direct structural centrality in tumor tissue.

MARCO^+^ macrophages showed different reorganization patterns, with 75.4% of metabolites (2,859 out of 3,793) maintaining original roles while 24.6% underwent transitions (Figure 5C, right panel). The most prominent transition was eigenvector-to-closeness (81 metabolites, 2.1%), where structurally central metabolites evolved toward enhanced connectivity. Degree-to-closeness transitions (60 metabolites, 1.6%) showed hub metabolites evolving toward efficient connectivity patterns.

We focused specifically on metabolites that transitioned from eigenvector-to-betweenness roles because they represent potential therapeutic targets with optimal selectivity profiles (Figure 5D). These metabolites evolved from highly connected, structurally central positions in normal tissue to critical bottleneck functions in tumor conditions. Subsystem analysis of these functionally critical metabolites revealed the specific metabolic pathways driving these architectural changes. In FAP^+^ fibroblasts, amino acid metabolism dominated the generation of new metabolic bottlenecks, including GSH, dihydrofolate, and adenylyl sulfate. L-dopaquinone emerged as a key transition metabolite contributing to seven distinct reactions in amino acid metabolism pathways. Hormone metabolism represented the second largest subsystem with six reactions, featuring GSH as a central coordinator. MARCO^+^ macrophages showed different patterns, with fatty acid metabolism representing the largest reorganized subsystem (35 reactions), featuring diverse acyl-CoA derivatives.

Analysis of extracellular metabolites identified 15 exchange metabolites with role transitions facilitating intercellular communication (Figure 5E). These included lipid signaling components (bile acid derivatives, prostaglandins), energy metabolites (CTP, UTP, lactose), amino acid trafficking molecules (alanine, glutamate), and antioxidant sharing mechanisms (GSH). The prominence of amino acid exchange metabolites directly corresponds to the extensive upregulation of amino acid metabolism observed in both FAP^+^fibroblasts and MARCO^+^ macrophages, while nucleotide exchange aligns with the enhanced nucleotide biosynthesis specifically in MARCO^+^ macrophages. Additionally, the presence of prostaglandin and bile acid derivatives among exchange metabolites reinforces our centrality analysis findings that these lipid mediators occupy critical network positions in tumor-associated stromal cells, suggesting these extracellular transitions coordinate the metabolic division of labor between the two populations.

### 2.7 Multifractal analysis reveals tumor-normal separation through geometric network properties

The metabolic adaptations identified through flux analysis and network topology may represent either isolated cellular responses or coordinated architectural changes emerging from TME interactions. In contrast, the multifractal and differential (e.g., Ollivier-Ricci curvature) geometric analyses can quantify the multiscale and global organizational principles. These analyses can identify whether tumor-associated cell types adopt similar geometric strategies for metabolic optimization.

Multifractal topological analysis reveals how metabolic networks are organized hierarchically from highly connected metabolic hubs that participate in many reactions (such as ATP and NADH), to specialized metabolites involved in only a few pathways. More precisely, the multifractal topological analysis investigates the statistical scaling behavior of the probability measure of covering the network with boxes of increasing sizes and estimates two main multifractal metrics: (i) the generalized fractal dimension *D*(*q*), which estimates the spectrum of fractal dimensions characterizing the network, and (ii) the multifractal spectrum *f* (*ω*), which provides information about the distribution of the Lipschitz-Holder scaling exponents (*ω*) observed across multiple subnetwork scales. Hence, multifractal topological analysis characterizes the hierarchical organization, the degree of heterogeneity, and the degree of complexity of networks by measuring how the scaling properties of nodes vary across different scales. Furthermore, this analysis can infer the generating rules governing how the networks were created.

The multifractal spectrum *f* (*ω*) (Figure 6A) captures three key geometric properties: complexity (*ω*_0_, the peak of the multifractal spectrum reflecting the dominant fractal dimension corresponding to the overall network intricacy), heterogeneity (spectrum width indicating diversity of structural elements), and asymmetry (left-right spectrum ratio distinguishing hub-dominated versus distributed network architectures). By examining how the scaling behavior of the density of metabolic connections expands with increasing box sizes (distances from each node), the multifractal spectrum in Figure 6A shows that the normal and tumor cells differ in the degree of asymmetry (see the right hand side tip of the multifractal spectra), degree of heterogeneity (see the range of Lipschitz-Holder exponents) and the complexity (the dominant [maximum] *ω*_0_ has the tendency to shift towards right from normal to tumor tissue). The right-shift tendency of *ω*_0_ and the increasing variability of the right hand side tip of the multifractal spectra indicate more chaotic dynamics corresponding to tumor condition.

**Fig. 6:**
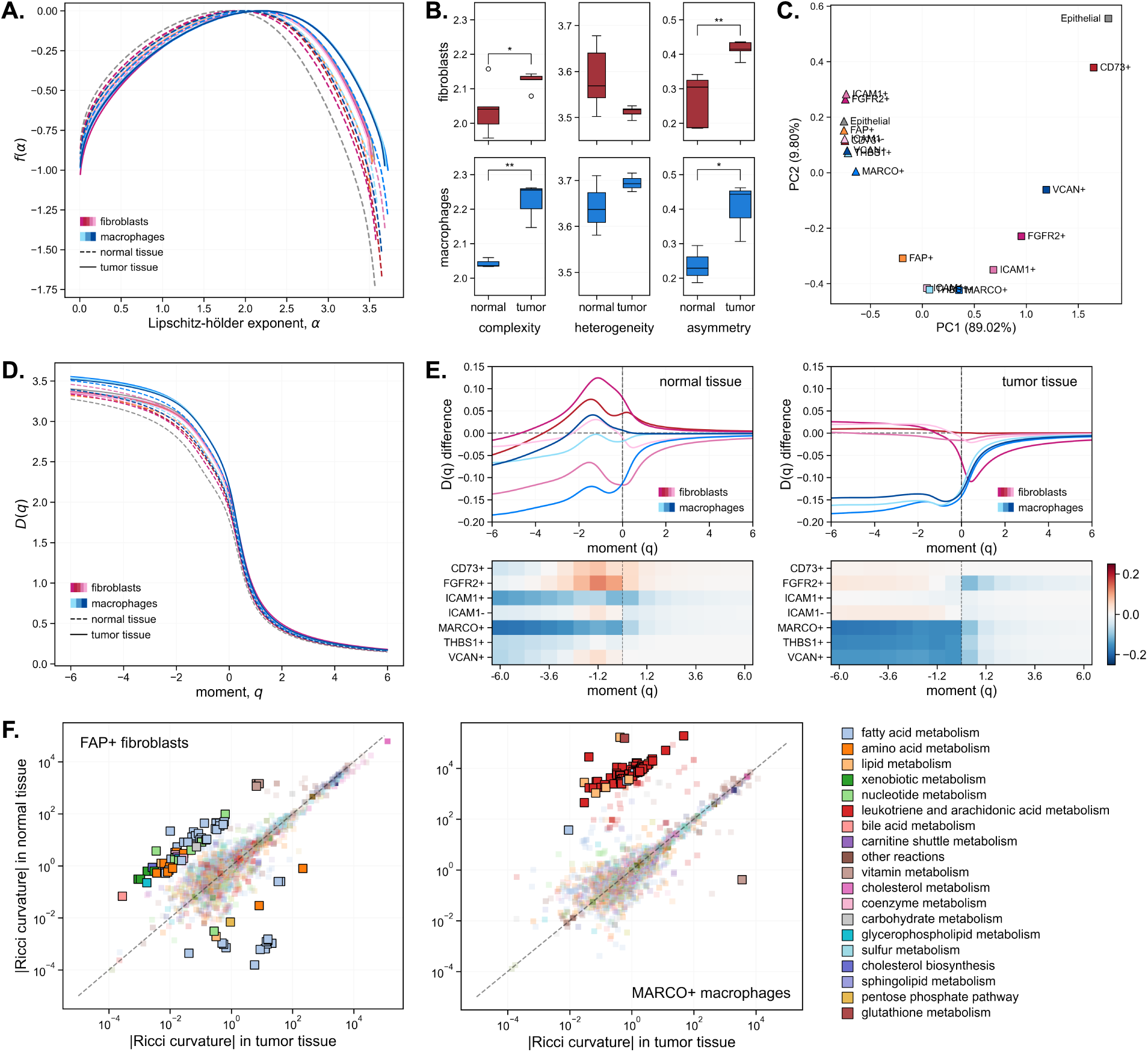
Geometric analysis reveals convergent evolution toward tumor-supporting network reorganization. **A.** Multifractal spectra *f* (*α*) characterizing network complexity across multiple scales, with complexity (*α*_0_), heterogeneity (spectrum width), and asymmetry (left-right ratio) properties extracted for each cell type. **B.** Quantification of geometric properties showing complexity, heterogeneity, and asymmetry values across cell types and tissue conditions. Statistical significance between tumor and normal conditions is indicated by asterisks (∗ *p <* 0.05, ∗∗ *p <* 0.01, ∗ ∗ ∗ *p <* 0.001). **C.** PCA of multifractal parameters separating tumor from normal phenotypes based on highest (*α*) values. **D.** Generalized fractal dimension *D*(*q*) curves across moment orders revealing scaling properties of metabolic network organization. **E.** Geometric distance analysis from FAP^+^ fibroblasts to other cell types in normal tissue (left) and tumor tissue (right), showing convergence patterns and ICAM1^+^ telocyte similarity to FAP^+^ fibroblasts in tumor conditions. **F.** Ollivier-Ricci curvature changes by metabolic subsystem for FAP^+^ fibroblasts (left) and MARCO^+^ macrophages (right), identifying pathways with most dramatic geometric reorganization in tumor conditions.

Quantification of these geometric properties revealed substantial differences between cell lineages and tissue conditions (Figure 6B). Complexity values demonstrated that tumor phenotypes maintain significantly higher network intricacy than their normal counterparts, suggesting adoption of more sophisticated metabolic architectures to adapt to TME conditions. Although not statistically significant between normal and tumor tissues, heterogeneity values showed that while macrophages tend to increase structural diversity in tumor conditions, fibroblasts became less heterogeneous. Asymmetry values revealed that tumorassociated stromal cells became significantly more asymmetrical, with the multifractal spectrum skewed toward sparsely connected regions. This suggests a more distributed organization and a higher likelihood of rare topological patterns.

We performed PCA on pairwise distance matrices derived from the highest *ω* values, corresponding to the most heterogeneous and sparsely connected regions of the metabolic networks. PCA revealed the first clear separation between tumor and normal tissue conditions across all cell types (Figure 6C). This result demonstrates that while individual geometric properties and single-scale network metrics failed to distinguish tumor from normal phenotypes, the multifractal approach successfully captures the subtle but coordinated multiscale architectural changes that characterize metabolic network reorganization in the TME.

Analysis of the generalized fractal dimension *D*(*q*) across different moment orders provides deeper insight into the multiscale properties that drive the observed separation in the multifractal spectrum (Figure 6D). To examine how these geometric changes affect relationships between cell types, we analyzed distances between FAP^+^ fibroblasts and other cell types using the generalized fractal dimension curves for different moment orders (Figure 6E). This distance metric captures differences in scaling behavior across the cell subtypes. Tumor conditions induced distinct reorganization patterns: ICAM1^+^ cells achieved near-complete geometric identity with FAP^+^ fibroblasts across all scaling regimes, maintaining near-zero distances from sparse to dense regions. In contrast, FGFR2^+^ fibroblasts showed increased divergence in highly connected areas in tumor conditions. At *q* ↑ 4, all cell types converge to similar values, indicating that the most highly connected network hubs follow universal scaling laws independent of cell type, likely reflecting shared metabolic constraints where currency metabolites and essential biosynthetic pathways dominate cellular organization.

Curvature analysis identified specific metabolic subsystems that undergo the significant changes in tumor conditions (Figure 6F). Specifically, the Ollivier-Ricci curvature metric quantifies the information flow throughout the network. The magnitude of the Ollivier-Ricci curvature changes identifies the pathways undergoing the most dramatic geometric reorganization. Points above the diagonal indicate reactions with higher Ollivier-Ricci curvature magnitudes in normal tissue, while points below the diagonal show higher magnitudes in tumor tissue, with the largest deviations representing the most significant pathway-level geometric adaptations. FAP^+^ fibroblasts showed the most pronounced curvature changes in fatty acid metabolism, amino acid metabolism, and xenobiotic metabolism, indicating fundamental geometric reorganization of these pathways in response to tumor conditions (Figure 6F, left panel). MARCO^+^ macrophages demonstrated different patterns of geometric reorganization, with the most significant curvature changes occurring in leukotriene and arachidonic acid metabolism (Figure 6F, right panel).

## 3 Discussion

Our comprehensive multi-scale modeling approach revealed multiple layers of tumor-associated vulnerabilities spanning flux upregulation, network topology reorganization, and geometric architectural changes to the metabolic network. FAP^+^ fibroblasts and MARCO^+^ macrophages exhibit extensive metabolic reprogramming with emphasis on carnitine shuttle metabolism, amino acid production, and redox detoxification pathways. Network topology analysis revealed metabolites undergoing centrality role transitions, particularly those shifting to critical bottlenecks. Geometric analysis identified pathway-specific reorganization patterns, with FAP^+^ fibroblasts showing dramatic curvature changes in fatty acid and xenobiotic metabolism, and MARCO^+^ macrophages exhibiting geometric optimization of inflammatory lipid mediator synthesis.

Traditional flux balance analysis assumes cellular metabolism optimizes for a single objective, typically biomass production. However, stromal cells in the TME face complex selective pressures requiring optimization across multiple competing objectives [38, 39]. Recent studies have shown that CAFs prioritize lactate production and collagen synthesis over their own proliferation [40, 41], while TAMs balance inflammatory mediator production with tissue remodeling functions [42, 43]. We employed random flux sampling to capture the full spectrum of feasible metabolic states, revealing that tumor FAP^+^ fibroblasts and MARCO^+^macrophages sacrifice growth capacity to support tumor progression. This metabolic altruism aligns with evidence from pancreatic cancer where CAFs undergo autophagy to provide amino acids to tumor cells [44], and extends observations from breast cancer where stromal cells adopt catabolic metabolism to fuel anabolic tumor growth [45].

The extensive upregulation of carnitine shuttle metabolism in FAP^+^ fibroblasts and MARCO^+^ macrophages indicates coordinated fatty acid oxidation programs serving dual purposes: generating energy for activated phenotypes while clearing lipid metabolites that could impair tumor cell function [46]. We also find divergent metabolic strategies—with FAP^+^ fibroblasts specializing in amino acid metabolism while downregulating nucleotide biosynthesis. This result parallels findings in pancreatic cancer where CAFs similarly upregulate amino acid catabolism [47]. Conversely, MARCO^+^ macrophages adopted cancer-like biosynthetic programs, consistent with studies showing TAMs exhibit enhanced purine metabolism [48].

Our systematic knockout analysis revealed metabolic vulnerabilities with direct clinical relevance and highlighted the complexity of translating computational predictions to therapeutic strategies. The exceptional tumor selectivity of MCCC1/MCCC2 (3-methylcrotonyl-CoA carboxylase) and the broader pattern of amino acid catabolism dependencies (BCAT2, PCCA, PCCB) validate our hypothesis that metabolic reprogramming creates shared vulnerabilities across tumor-associated populations. These enzymes catalyze essential steps in branched-chain amino acid degradation, and their selective essentiality aligns with evidence that BCAA metabolism is fundamentally rewired in CRC [49]. The convergence of multiple amino acid catabolism enzymes (including MCEE, MMUT in propionate metabolism and HGD and DHTKD1 in lysine degradation) as top-ranking candidates underscores the central importance of amino acid supply networks in sustaining the tumor ecosystem.

Our metabolite role transition analysis represents a significant advance in identifying context-specific therapeutic targets. By tracking how metabolites shift between functional roles in normal and tumor conditions, we identified metabolic control points that emerge specifically in the TME. In FAP^+^ fibroblasts, the prominence of glutathione and dihydrofolate among transition metabolites aligns with evidence that CAFs undergo oxidative stress requiring enhanced antioxidant capacity [50]. Studies targeting glutathione metabolism in CAFs show potential for overcoming therapeutic resistance [51].

MARCO^+^ macrophages showed distinct vulnerabilities centered on fatty acid metabolism, with multiple acyl-CoA species transitioning to bottleneck roles. This comprehensive reorganization reflects the macrophages’ evolution into specialized lipid processors, consistent with lipidomic analyses showing TAMs accumulate specific lipid species promoting immunosuppression [52]. The identification of extracellular metabolites undergoing role transitions provides direct evidence for metabolic communication networks. Prostaglandin derivatives emerged as central mediators of this cell-cell metabolic communication, with recent clinical trials targeting prostaglandin synthesis showing promise in CRC prevention [53].

The application of multifractal analysis and Ollivier-Ricci curvature to metabolic networks represents a methodological innovation that revealed previously hidden patterns of metabolic coordination and similarities across cell subtypes. While conventional network metrics failed to distinguish tumor from normal phenotypes, geometric analysis successfully captured coordinated architectural changes characterizing metabolic adaptation in the TME. The increased network complexity and asymmetry in tumor-associated populations suggests adoption of sophisticated metabolic architectures capable of navigating the challenging TME. Our finding that ICAM1^+^ telocytes achieve near-complete geometric convergence with FAP^+^ fibroblasts provides mechanistic insights into stromal cell differentiation, supporting RNA-velocity analysis of Qi et al [17]. The scale-dependent geometric reorganization suggests different precursor populations follow distinct trajectories to the cancer-associated fibroblast state.

Our findings reveal that metabolic adaptation represents a coordinated evolutionary process rather than random changes. The metabolic division of labor between FAP^+^ fibroblasts and MARCO^+^ macrophages resembles metabolic syntrophy observed in microbial communities [54]. The prominence of carnitine shuttle metabolism extends recent observations that fatty acid oxidation fuels immunosuppressive functions [46]. Prostaglandin and bile acid metabolism emerged as metabolic highways of stromal crosstalk. Beyond established roles in inflammation and tumor promotion [55], our network analysis reveals these metabolites occupy topologically critical positions specifically in tumor-associated populations.

To assess the translational potential of our computational findings, we surveyed existing preclinical and clinical evidence for the metabolic targets and pathways identified through our integrated analysis (Table 1). Many of our predictions align with active areas of therapeutic development, providing independent validation that our ecosystem-level approach identifies biologically and clinically relevant vulnerabilities. Most notably, MAOB has recently emerged as a therapeutic target in colorectal cancer, with MAO inhibitors currently being investigated for their anti-tumor efficacy [56, 57]. Similarly, our network analysis positioning prostaglandins as critical hubs in FAP^+^ fibroblasts provides mechanistic insight into decades of epidemiological and clinical trial evidence supporting COX-2 inhibitors for CRC prevention [58, 59]. The convergence of our computational predictions with these established therapeutic programs demonstrates that metabolic network topology can identify clinically actionable targets. Furthermore, our novel findings, including the role transitions of specific metabolites and the geometric reorganization patterns, point toward previously unrecognized intervention opportunities.

**Table 1:** Translational evidence supporting computationally identified metabolic vulnerabilities. Computational findings from our integrated analysis are matched with existing therapeutic strategies and development programs. Development stages reflect highest reported level of clinical evaluation for cancer indications.

| Computational Finding | Therapeutic Target | Intervention | Development Stage | Ref. |
| --- | --- | --- | --- | --- |
| *Gene knockout vulnerabilities (from Figure 2)* |  |  |  |  |
| BCAT2 gene essentiality in tumor cells | BCAA catabolism | BCAT inhibitors; dietary BCAA restriction | Preclinical | [60, 61] |
| MCCC1/2 tumor-selective lethality | Leucine degradation | — | Emerging target | [62] |
| MAOB gene survival association | Monoamine oxidase B | Rasagiline; Selegiline; Deprenyl | Preclinical (repurposing) | [56, 57, 63] |
| PCCA/PCCB dependency | Propionate metabolism | Metabolic modulators | Preclinical | [64] |
| TYMS gene essentiality in tumor cells | Thymidylate synthase | 5-Fluorouracil; Capecitabine | Approved (standard of care) | [65, 66] |
| *Metabolic specialization (from Figures 1, 5)* |  |  |  |  |
| FAP<sup>+</sup> fibroblast targeting | Fibroblast activation protein | Anti-FAP CAR-T; FAPI radioligands; Simlukafusp alfa | Phase I/II | [67–69] |
| Carnitine shuttle activity | CPT1A/CPT2 | Etomoxir; Perhexiline | Preclinical | [70–72] |
| *Network topology bottlenecks (from Figures 4, 5)* |  |  |  |  |
| Prostaglandin centrality | COX-2/PGE2 | Celecoxib; Aspirin | Phase III (prevention); Phase II (adjuvant) | [58, 59, 73] |
| GSH role transitions | Glutathione synthesis | BSO (buthionine sulfoximine) | Phase I | [74, 75] |
| Dihydrofolate transitions | DHFR | Methotrexate; Pralatrexate | Approved | [76] |
| Bile acid hub centrality in FAP<sup>+</sup> fibroblasts | FXR/TGR5 signaling | Obeticholic acid (OCA); Fexaramine D | Approved (PBC); Preclinical (CRC) | [77–79] |
| *Geometric network reorganization (from Figure 6)* |  |  |  |  |
| Fatty acid curvature changes in FAP<sup>+</sup> fibroblasts | FASN; ACC | TVB-2640 | Phase II | [80] |
| Leukotriene/arachidonic acid curvature changes in MARCO<sup>+</sup> macrophages | 5-LOX/LTA4H | Zileuton | Approved (asthma); Pre-clinical (cancer) | [81, 82] |
| *Ecosystem-level strategies* |  |  |  |  |
| Metabolic exchange disruption | MCT1 (lactate transport) | AZD3965 | Phase I (completed) | [83, 84] |

*Abbreviations:* BCAA, branched-chain amino acids; AA, amino acid; GSH, glutathione; BSO, buthionine sulfoximine; FAO, fatty acid oxidation; DHFR, dihydrofolate reductase; FXR, farnesoid X receptor; 5-LOX, 5-lipoxygenase.

We generated several testable hypotheses for experimental validation. First, we hypothesize that metabolic division of labor between FAP^+^ fibroblasts and MARCO^+^ macrophages suggests these populations coordinate through metabolite exchange mechanisms that could be mapped using organoid co-culture systems with metabolic tracing. Second, our identification of metabolites undergoing role transitions to bottleneck positions suggests that targeting these network control points may provide therapeutic advantages over conventional single-pathway approaches. Third, we propose that the geometric reorganization patterns we observed indicate that metabolic network architecture undergoes coordinated changes in tumor conditions, which could be further characterized using perturbation experiments combined with network analysis. Integration of such experimental validation with expanded computational models will enhance predictive accuracy and reveal additional therapeutic vulnerabilities in the colorectal cancer ecosystem.

Our study introduces several methodological innovations that advance the field of cancer metabolism beyond traditional metabolic engineering approaches. The application of multifractal analysis and Ollivier-Ricci curvature to networks reconstructed from GEMs provides new tools for understanding metabolic organization at multiple scales. However, some limitations warrant consideration for future improvements. Our reliance on transcriptomic data for model constraints, while standard practice, may not fully capture post-translational regulation of metabolic enzymes. Integration with proteomic and metabolomic data could improve model accuracy significantly. Future studies incorporating spatial metabolomics and singlecell metabolic profiling will provide higher resolution views of metabolic organization and validate our computational predictions.

## 4 Conclusion

Our systems biology approach reveals that metabolic reprogramming in the CRC microenvironment represents a coordinated evolutionary process where FAP^+^ fibroblasts and MARCO^+^ macrophages function as metabolic cooperators, sacrificing individual fitness for collective tumor support. This cooperation creates an extended ecosystem where tumor properties emerge from multicellular metabolic networks rather than cancer cells alone.

By integrating analytical scales from reaction fluxes to network topology and geometric organization, we bridge the critical gap between molecular mechanisms and systems-level behavior in cancer metabolism research. Our findings demonstrate that minor changes at the reaction level can drive dramatic network reorganization, explaining why single-pathway targeting often fails [85].

The identification of shared metabolic dependencies, critical bottlenecks, and geometric reorganization patterns provides a road map for targeting the metabolic ecosystem rather than individual cell types. As precision oncology advances, metabolic network architecture should guide therapeutic decisions. The success of immunotherapy has demonstrated the power of targeting cellular interactions rather than cancer cells directly [86]. Similarly, targeting metabolic interactions and network properties may overcome the limitations of current metabolic therapies that focus on individual enzymes or pathways [87]. Our findings reveal that metabolic adaptation in cancer emerges through coordinated, ecosystem-level reorganization of interacting cell populations, outlining a general framework for therapeutic strategies that target cooperative metabolic networks rather than isolated pathways. While we demonstrate this ecosystem-targeting strategy in CRC, the principles of metabolic cooperation between tumor-associated fibroblasts and macrophages likely extend across other solid tumors, and our scalable framework enables systematic identification of these vulnerabilities in any cancer type with available data. This work has the potential to transform how we approach therapeutic resistance across oncology.

## 5 Methods

### 5.1 ScRNA-seq data acquisition and processing

We used a publicly available single-cell RNA-seq dataset of colorectal cancer comprising 54,103 cells across tumor and tumor-adjacent normal tissues [17]. The original study’s cell type annotations were retained to define biologically distinct populations. In particular, annotated cell types included stromal fibroblasts (subtypes characterized by FAP, FGFR2 or NT5E/CD73 expression) and telocytes (ICAM1^+^ or ICAM1*^→^* subclusters), immune macrophage subtypes (MARCO^+^, THBS1^+^, VCAN^+^), as well as malignant epithelial cells in both tissues. Cell clusters with fewer than 50 cells per tissue type were excluded to ensure robust expression profiles. Pseudobulk expression profiles were generated for each cell type by averaging gene expression values across all cells within each cluster and tissue type, mitigating technical noise while preserving cell type-specific transcriptional signatures[88]. Data processing were performed using R (v4.0+) with Seurat package[89].

### 5.2 Genome-scale metabolic model preparation and context-specific model generation

The Human-GEM model (v1.19)[25] was used as the foundation for context-specific model generation. The model was extended with a reactive oxygen species (ROS) module[90] to capture redox metabolism in the tumor microenvironment. Integration was performed by matching metabolites using KEGG identifiers and establishing consistent reaction nomenclature, resulting in 13,213 reactions and 8,621 metabolites.

Context-specific models for each cell type and tissue were derived using the transcriptome-driven ftINIT algorithm [25]. Briefly, the algorithm maps the gene expression values to reaction scores using gene–protein–reaction (GPR) rules: reactions with an OR relationship used the maximum gene expression, while those with an AND relationship used the minimum [27]. The algorithm retained all high-activity reactions (*>*50th percentile), then and added lower-activity reactions required to fulfill 57 predefined essential metabolic tasks [91]. We did not impose any external constraints on nutrient uptake rates as these parameters are highly variable in the tumor microenvironment and poorly characterized for human tumors in situ. Each model was verified for biomass production.

### 5.3 Random flux sampling

Random flux sampling was performed using Riemannian Hamiltonian Monte Carlo (RHMC)[92] implemented in the COBRA Toolbox[93]. Each model was constrained to maintain ↑50% of maximum biomass production to ensure physiologically relevant flux distributions, with 2,000 samples generated per model. This ↑50% threshold was selected to exclude metabolically compromised or dying cells while allowing exploration of non-proliferative states.

Due to the high-dimensional nature of genome-scale models, RHMC can generate samples that include thermodynamically inconsistent flux loops, particularly in sparsely constrained regions of the solution space. To eliminate thermodynamically infeasible loops inherent in random sampling, corner sampling[94] was performed using RAVEN Toolbox[95] to define feasible flux ranges by minimizing total flux. RHMC samples were filtered by setting reaction fluxes to zero when their mean values exceeded corner sampling ranges by *>*5 units, effectively removing futile cycles while preserving the statistical power of the sampling approach. All flux distributions were normalized to unit biomass production (growth rate = 1.0) before downstream analysis to ensure comparable flux magnitudes across different cell types and conditions.

### 5.4 Differential reaction activity analysis

Differential reaction activity between tumor and normal tissues was assessed using four complementary statistical criteria: (i) Mann–Whitney U test (FDR *<* 0.01), (ii) absolute log_2_ fold-change ↑ 2.0, (iii) Cohen’s *d* effect size ↑ 0.5, and (iv) Wasserstein distance ↑ 5.0. Given the large sample sizes and multiple cell types analyzed, statistical significance was achieved for numerous small-magnitude differences. The stringent thresholds, particularly the Wasserstein distance criterion, were implemented to focus on biologically meaningful large-magnitude changes representing primary metabolic adaptations rather than minor fluctuations. Pathway-level analysis was performed by aggregating results according to metabolic subsystem annotations in Human-GEM after aggregating similar subsystems.

### 5.5 in silico gene knockout analysis

To validate and prioritize identified metabolic targets, we performed systematic computational gene knockout analysis. Single gene deletion simulations were performed on each cell type-specific model using flux balance analysis with minimization of metabolic adjustment (MOMA) [96]. For each candidate gene, the corresponding reactions were constrained to zero flux, and the resulting impact on biomass production was calculated as knockout ratio (post-knockout biomass/wild-type biomass). Potential therapeutic targets were identified using dual-criteria filtering requiring significant tumor growth inhibition (mean tumor cell viability → 0.4) and minimal normal cell toxicity (mean normal cell viability ↑ 0.6). Identified targets were then prioritized based on their most active metabolic pathways, with genes affecting critical pathways (fatty acid metabolism, amino acid metabolism, nucleotide metabolism, ROS detoxification) ranked higher than those in less essential pathways. Within each pathway category, genes were further ordered by their effectiveness in suppressing tumor cell viability.

### 5.6 Target validation and survival correlation

Clinical validation of computationally identified metabolic targets was performed using survival analysis with the GSE17536 colorectal cancer dataset[97], which contains gene expression and clinical outcome data from CRC patients. All genes that passed the knockout analysis filtering criteria were subjected to KaplanMeier survival analysis for both overall survival (OS) and disease-free survival (DFS) using optimal cutpoint analysis.

### 5.7 Metabolic network graph construction

Directed metabolite-centric graphs were constructed from each cell type-specific model using the stoichiometric matrix and mean flux values from random sampling. For each reaction, metabolites with negative stoichiometric coefficients (substrates) were connected to those with positive coefficients (products) via directed edges. Edge weights corresponded to mean reaction fluxes, and only reactions with non-zero flux were included. Networks were filtered to retain the largest connected component containing the biomass metabolite, ensuring focus on active metabolic pathways[98].

### 5.8 Topological characterization

We computed over 90 network metrics encompassing: (i) basic structural properties including node and edge counts, network density; (ii) degree-based measures including centralization, heterogeneity, and distribution skewness; (iii) weight-based statistics capturing flux asymmetry and distribution moments; and (iv) structural node classifications identifying branch points, linear pathways, terminal nodes, sources, and sinks, using NetworkX package[99].

Nodes of each graph were classified by dominant centrality measures: degree centrality (normalized connection count), betweenness centrality (frequency on shortest paths between other nodes), closeness centrality (inverse of average shortest path distance), PageRank (recursive importance considering neighbor importance), load centrality (fraction of shortest paths passing through the node), and eigenvector centrality (importance based on connections to other important nodes). For directed networks, separate in-degree and out-degree centralities were computed.

### 5.9 Metabolite role transitions

Each metabolite’s centrality values were converted to percentile ranks within its respective network, and the metabolite was assigned to the functional role corresponding to its highest-ranking centrality measure. This yielded six metabolite categories: (i) highly connected hubs (dominant degree centrality), (ii) critical bottlenecks connecting distinct network modules (dominant betweenness centrality), (iii) highly accessible nodes with efficient connections throughout the network (dominant closeness centrality), (iv) prominent nodes linked to other topologically important metabolites (dominant PageRank), (v) high-traffic carriers of network flow (dominant load centrality), and (vi) structurally central nodes in the overall network architecture (dominant eigenvector centrality).

Metabolic reorganization was quantified by analyzing role transitions between tumor and normal networks using transition matrices, where each element represents the number of metabolites shifting from one functional role to another. Network-level comparisons employed k-means clustering on standardized topological metrics, with optimal cluster number determined by silhouette analysis. Principal component analysis of centrality matrices identified systematic patterns of metabolic control point reorganization across cell types and tissue conditions.

### 5.10 Multifractal geometric analysis

To determine whether identified metabolic differences in the tumor microenvironment represent coordinated adaptation, we characterized the global organizational principles of metabolic networks using geometric analysis.

Multifractal analysis was performed using node-based multifractal analysis (NMFA)[100] to characterize network complexity across multiple scales. For each node, the scaling relationship between mass distribution *M* (*r*) and radius *r* was calculated, yielding node-specific fractal dimensions. The partition function was computed across moment orders *q* ↓ [↔10, 10], and the multifractal spectrum f(*ω*) was derived using the approach described by Xiao et al. Key metrics extracted included complexity (*ω*_0_, spectrum peak position reflecting overall network intricacy), heterogeneity (spectrum width indicating diversity of structural elements), and asymmetry (left-right spectrum ratio distinguishing hub-dominated versus distributed network architectures). Pairwise network distances were calculated using Wasserstein distance on multifractal spectra, enabling quantitative geometric comparison between cell types and tissue conditions[100].

### 5.11 Curvature analysis

To quantify structural differences between normal and tumor metabolic networks, we applied Ollivier-Ricci curvature analysis. Networks were first simplified by combining parallel edges between identical metabolite pairs, summing reaction weights to represent cumulative interaction strength. Ollivier-Ricci curvature was calculated for each edge using the GraphRicciCurvature package [101] with idleness parameter *ω* = 0.5. We compared curvature magnitudes between corresponding reactions across conditions, calculating absolute log2 fold changes to identify the most structurally altered reactions, where tumor conditions induce the greatest topological changes in network structure.

### 5.12 Software and Code Availability

All analyses were performed using: MATLAB R2025b with COBRA Toolbox v3.0 and RAVEN Toolbox v2.9.0 for metabolic modeling; Python 3.12.4 with NumPy v1.26.4, pandas v2.2.2, SciPy v1.15.1, scikit-learn v1.5.2, and statsmodels v0.14.4 for data analysis; Matplotlib v3.9.2, Seaborn v0.13.2, and Plotly v5.24.1 for visualization; NetworkX v3.4.2, NetworKit v11.0, and GraphRicciCurvature v0.5.3.2 for network analysis. Custom scripts are available at: https://github.com/FinleyLabUSC/tme-network-topology.

## 6 Acknowledgements

The authors thank the members of the Finley Lab at USC for the deep scientific discussions and critical feedback. H.C. and S.D.F. disclose support for the research of this work from the USC Center for Computational Modeling of Cancer.

## 7 Author Contributions

H.C. conducted all metabolic modeling and computational analyses. P.B. and S.M. contributed to scientific discussions of results. S.D.F. was responsible for the scientific direction of this work. All authors contributed to manuscript writing and reviewed and provided feedback on the manuscript.

