## Supplementary Figures for "Metabolic Reprogramming and Therapeutic Vulnerabilities in the Tumor Microenvironment Revealed by Multi-scale Network Geometry"

This supplement provides additional analyses supporting the main findings of our study. **Figure S1** demonstrates that metabolic network architecture serves as a robust cellular fingerprint, with hierarchical clustering of binary reaction vectors showing strict separation by cell lineage (fibroblast vs. macrophage) rather than by tissue condition (tumor vs. normal). **Figure S2** presents detailed reaction-level flux changes across all metabolic subsystems, revealing the specific biochemical reactions underlying the pathway-level reprogramming patterns described in the main text, including exchange reaction patterns that suggest potential metabolic crosstalk mechanisms between tumor-associated cell populations. **Figure S3** shows that individual topological network metrics fail to distinguish tumor from normal conditions when analyzed in isolation, motivating our multifractal geometric approach that successfully captures coordinated architectural changes across multiple scales.

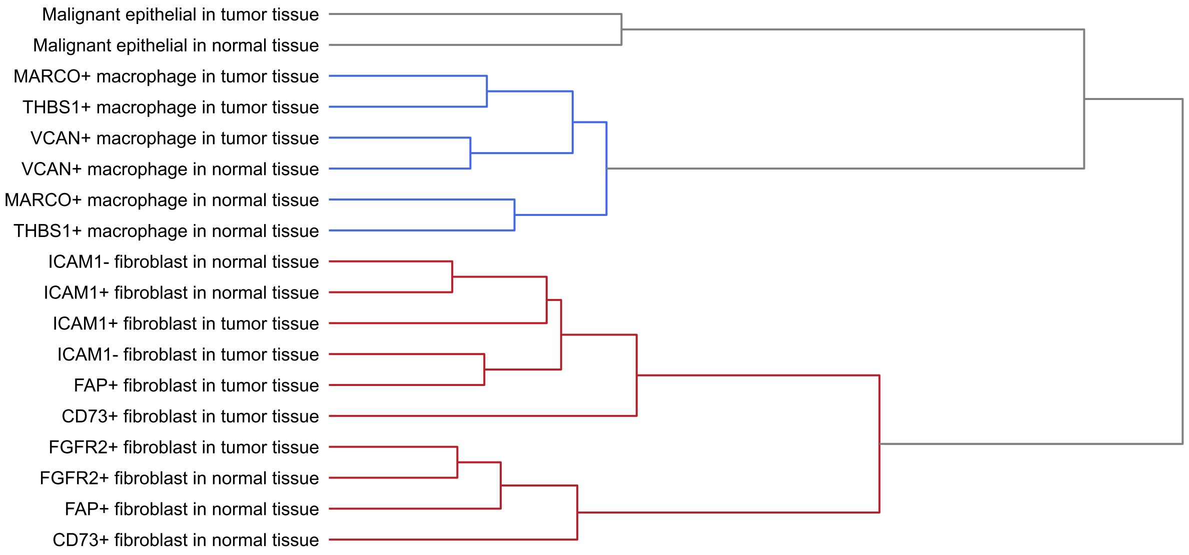


**Figure S1.** Hierarchical clustering of binary reaction vectors demonstrating separation between fibroblast and macrophage lineages by metabolic architecture.

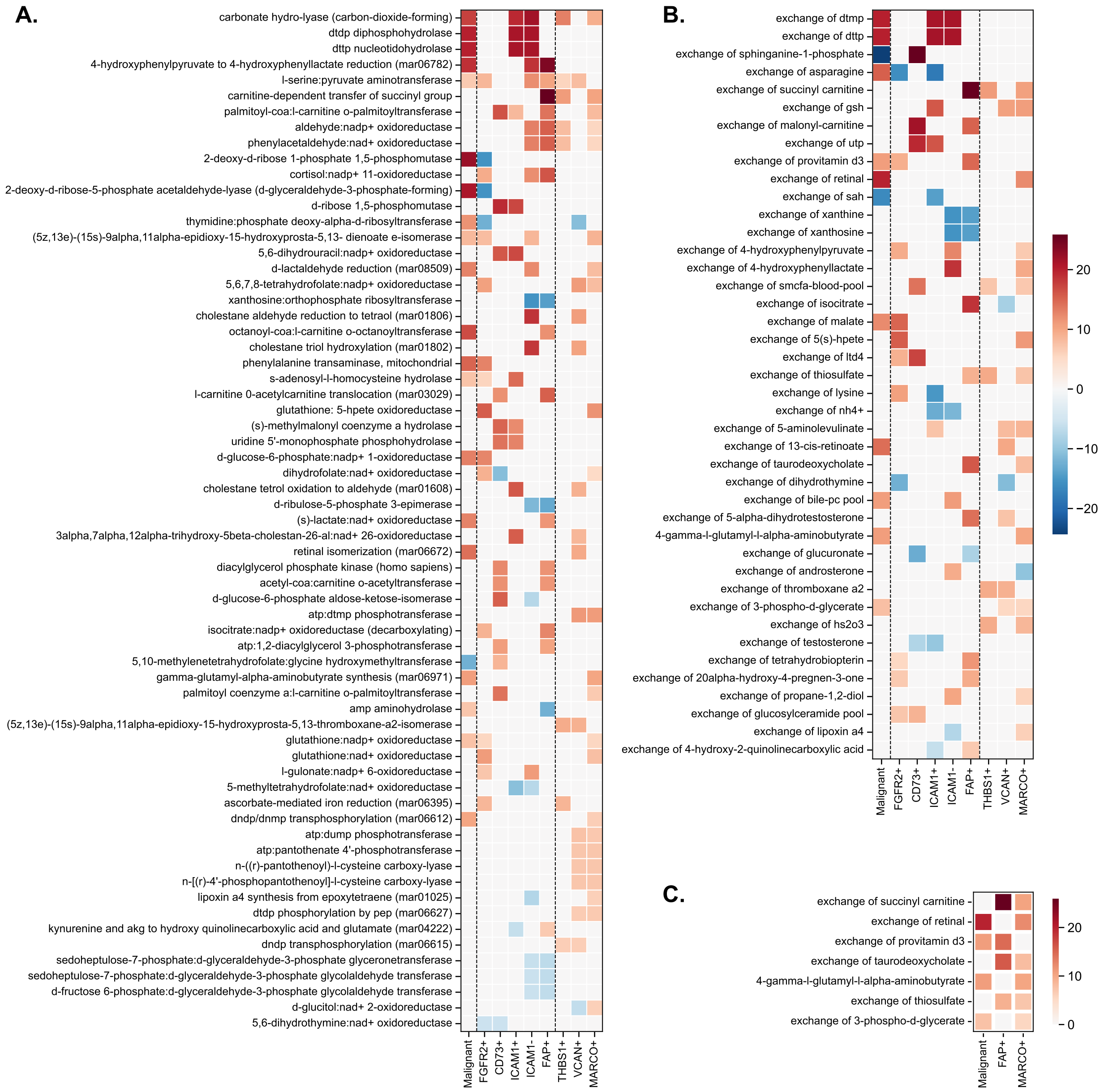


**Figure S2.** Detailed reaction-level analysis reveals complementary metabolic reprogramming patterns. **A.** Heatmap of individual reactions across all metabolic subsystems showing Wasserstein distance values (positive values indicate upregulation in tumor conditions) for all cell types, with focus on the most significantly altered reactions. **B.** Exchange reaction patterns showing differential uptake and secretion capabilities between cell types. **C.** Focused analysis of exchange reactions in malignant epithelial cells, $MARCO^{+}$ macrophages, and $FAP^{+}$ fibroblasts, highlighting metabolic dependencies and potential crosstalk mechanisms. Color scale represents Wasserstein distance with positive values indicating increased flux in tumor conditions.

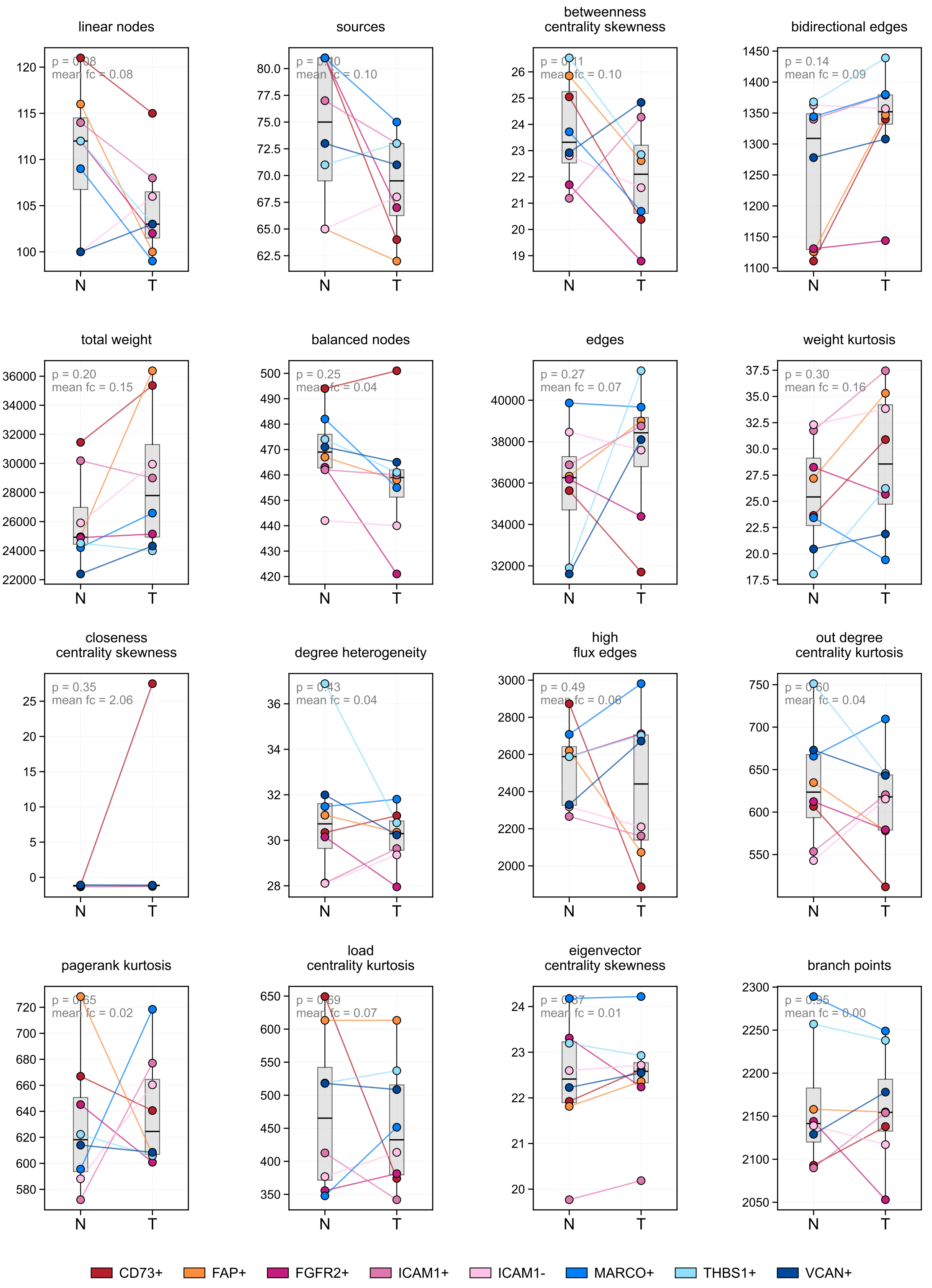


**Figure S3.** Individual topological features show no clear separation between tumor and normal conditions. Box plots comparing 16 selected topological metrics between normal (N) and tumor (T) metabolic networks across multiple cell subtypes.
